# Machine Learning for the Identification of Viral Attachment Machinery from Respiratory Virus Sequences

**DOI:** 10.1101/2022.01.25.477734

**Authors:** Stepan Demidkin, Maïa Shwarts, Arijit Chakravarty, Diane Joseph-McCarthy

## Abstract

At the outset of an emergent viral respiratory pandemic, sequence data is among the first molecular information available. As viral attachment machinery is a key target for therapeutic and prophylactic interventions, rapid identification of viral “spike” proteins from sequence can significantly accelerate the development of medical countermeasures. For five families of respiratory viruses, covering the vast majority of airborne and droplet-transmitted diseases, host cell entry is mediated by the binding of viral surface glycoproteins that interact with a host cell receptor. In this report it is shown that sequence data for an unknown virus belonging to one of the five families above provides sufficient information to identify the protein(s) responsible for viral attachment and to permit an assignment of viral family. Random forest models that take as input a set of respiratory viral sequences can classify the protein as “spike” vs. non-spike based on predicted secondary structure elements alone (with 97.8 % correctly classified) or in combination with N-glycosylation related features (with 98.1 % correctly classified). In addition, a Random Forest model developed using the same dataset and only secondary structural elements was able to predict the respiratory virus family of each protein sequence correctly 89.0 % of the time. Models were validated through 10-fold cross-validation as well as bootstrapping. Surprisingly, we showed that secondary structural element and N-glycosylation features were sufficient for model generation. The ability to rapidly identify viral attachment machinery directly from sequence data holds the potential to accelerate the design of medical countermeasures for future pandemics.

## Introduction

The COVID-19 pandemic has underscored the importance of an effective response for emerging viral pathogens that is focused on the rapid deployment of molecular testing and medical countermeasures. Our experiences with the current pandemic have highlighted the vulnerability of the global healthcare infrastructure to respiratory pathogens that, like SARS-CoV-2, are capable of long-range airborne spread via aerosolized particles [1]. In contrast to other pathogens, the window for effective intervention to avert a pandemic resulting from a newly emergent respiratory virus may be very short. Thus, the speed with which molecular diagnostics, therapeutics, and vaccines can be deployed are critical determinants of our ability to contain an outbreak.

The viral attachment machinery (the set of proteins responsible for host cell attachment and cell entry) has served as a historically important focus for the development of molecular tests (for example for influenza [2] and SARS-CoV-2 [3, 4]) as well as medical countermeasures such as vaccines [5–7]. Thus, the accurate and efficient identification of the viral attachment machinery is a critical first step in the design and deployment of biomedical countermeasures. It had been observed for coronaviruses in 2012 (pre-COVID-19) that the tertiary structure of the spike protein is not conserved but that the secondary structure topology is conserved [8]. It was subsequently also noted that the pattern of N-linked glycosylation is highly conserved and may play a role in immune evasion [9].

Automated function prediction (AFP) of novel proteins is a mature field (see [10–13] for reviews). A number of groups have used approaches that leverage structure-based homology, focusing either on the full three-dimensional (3D) protein structure, or on the identification of 3D structural motifs (see, for example, [14–17]). However, 3D structure alone is often insufficient for functional annotation, as proteins possessing similar global structures can perform very different biological functions (for example, [18]). Computational structural alignment methods, although first pioneered in the 1960s, typically have accuracies on the order of ~90% [19] and at least in the case of coronaviruses as described above the 3D structure is not conserved. Furthermore, 3D structural motifs for viral attachment proteins are often optimized for specifically for enzymes and are not readily able to identify viral attachment machinery. As an alternative, AFP from DNA sequences relies on sequence homology [20–22], or the identification of sequence motifs [23, 24]. A potential weakness of this approach is that novel viruses with low sequence homology to pre-existing pathogens may prove less tractable to homology-based approaches. As a further consideration, during the early days of an emerging pandemic, steps such as multiple sequence alignment, phylogeny reconstruction and 3D structure prediction can add weeks to the timeline for response. An accurate ML model may be able to pinpoint the target within seconds.

With respect to preparedness for potential future pandemics, tools that can aid in the rapid deployment of therapeutic and vaccine countermeasures are clearly needed. Specifically, for viral pathogens originating from the most prevalent respiratory virus families, which are key pathogens of concern, intervening at the localized emergence stage may prevent the transition to a full-blown pandemic. Based on the earlier cited observations, we hypothesized it may be possible to develop a machine learning (ML) model based on predicted secondary structure elements and N-glycosylation features alone capable of identifying viral attachment machinery (the “spike” protein or its equivalent) from an unknown respiratory virus sequence. More generally, we also sought to gain a further understanding of the structural features that may distinguish viral attachment machinery proteins with a view toward elucidation of key structure-function relationships.

## Methods

### Virus families, viral sequences, and “spike” proteins

Five families of respiratory viruses were included in this study: Coronaviridae, Paramyxoviridae, Pneumoviridae, Adenoviridae, and Orthomyxoviridae. Each of the viruses within these families has a protein responsible for viral attachment and host cell entry, which will be referred to herein as the “spike” protein (see Fig. 1A). For Coronaviruses, it is the Spike S Glycoprotein which is aptly named because it projects from the surface of the virion (Fig. 1B) as do the other “spike” proteins. Note that for Influenza Virus A within the Orthomyxoviridae family, we selected Hemagglutinin as the equivalent of the “spike” although Neuraminidase is a second antigenic determinant. A total of 39 sequences (ranging from 4 to 12 for each virus family) encoding 316 proteins were utilized (see Table 1). Specifically, we included 7 Coronaviridae sequences representing 7 viruses, 4 Paramyxoviridae sequences representing 4 viruses, 12 Pneumoviridae sequences representing 2 viruses, 8 Adenoviridae sequences representing 1 virus, and 8 Orthomyxoviridea sequences representing 1 virus.

**FIG 1.**
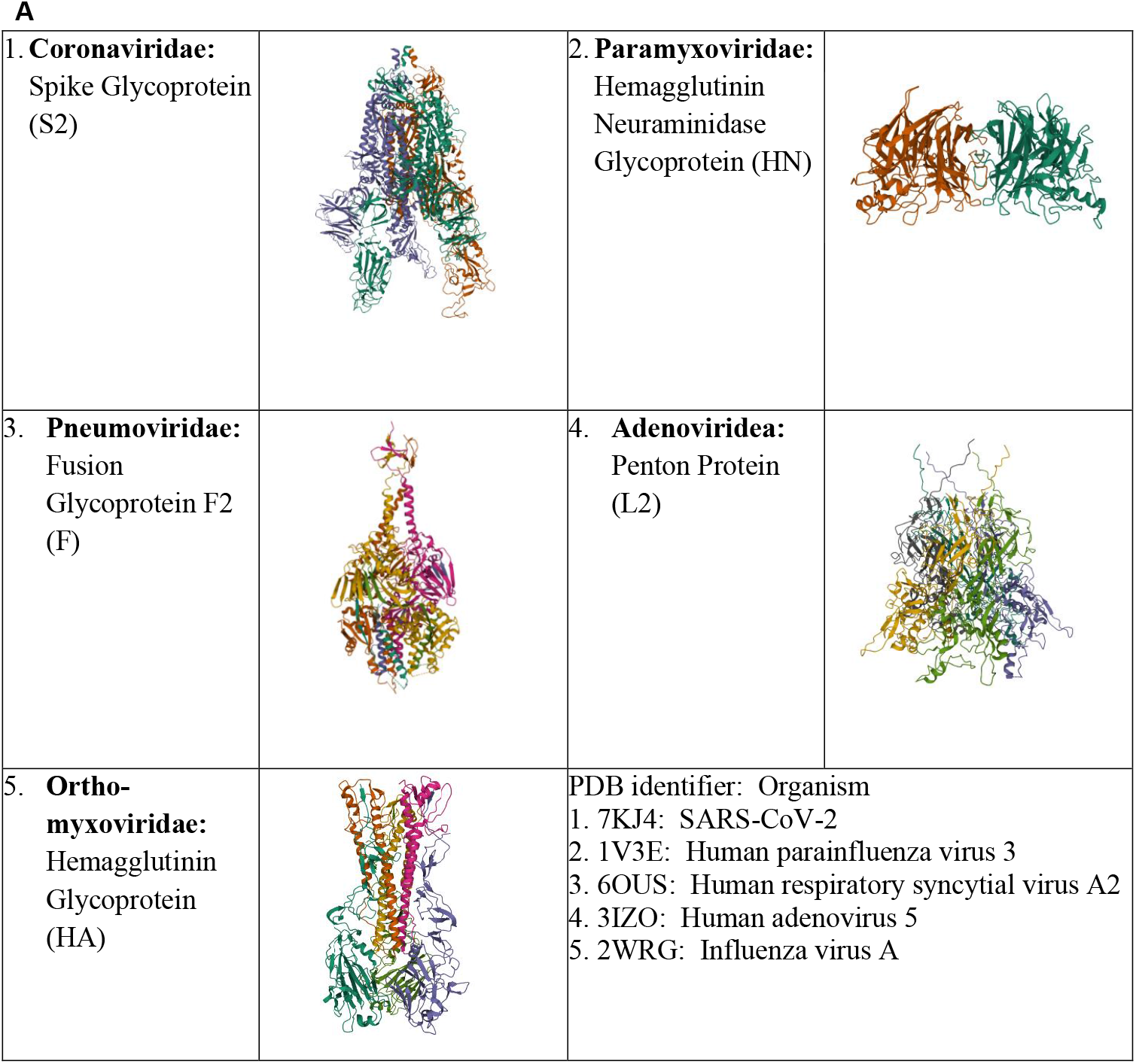
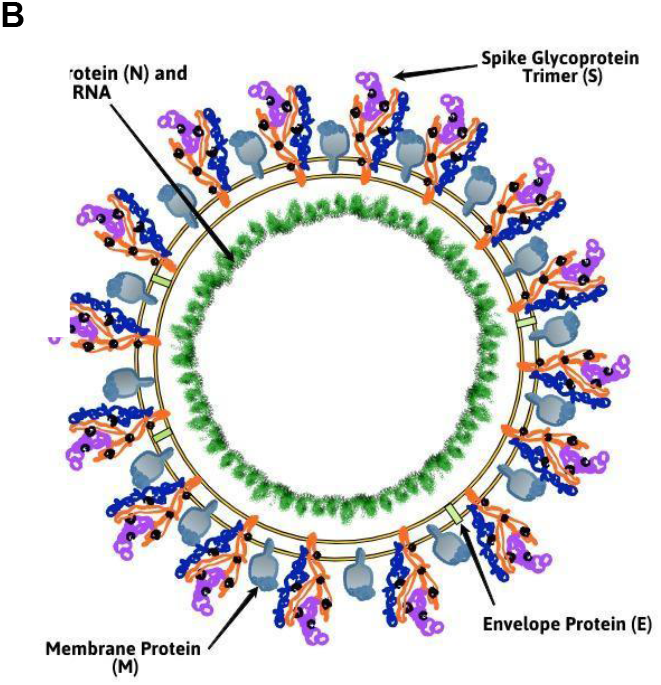
Five families of respiratory viruses and their “spike” proteins. In (A) the identity and representative structure of the “spike” protein (gene name given in parentheses) is shown for each of the virus families studied. PDB identifiers for structures 1-5 are also listed with the corresponding virus indicated. Shown in (B) is a schematic of the coronavirus SARS-CoV-2 structure indicating the prominence of the spike.

**FIG 2.**
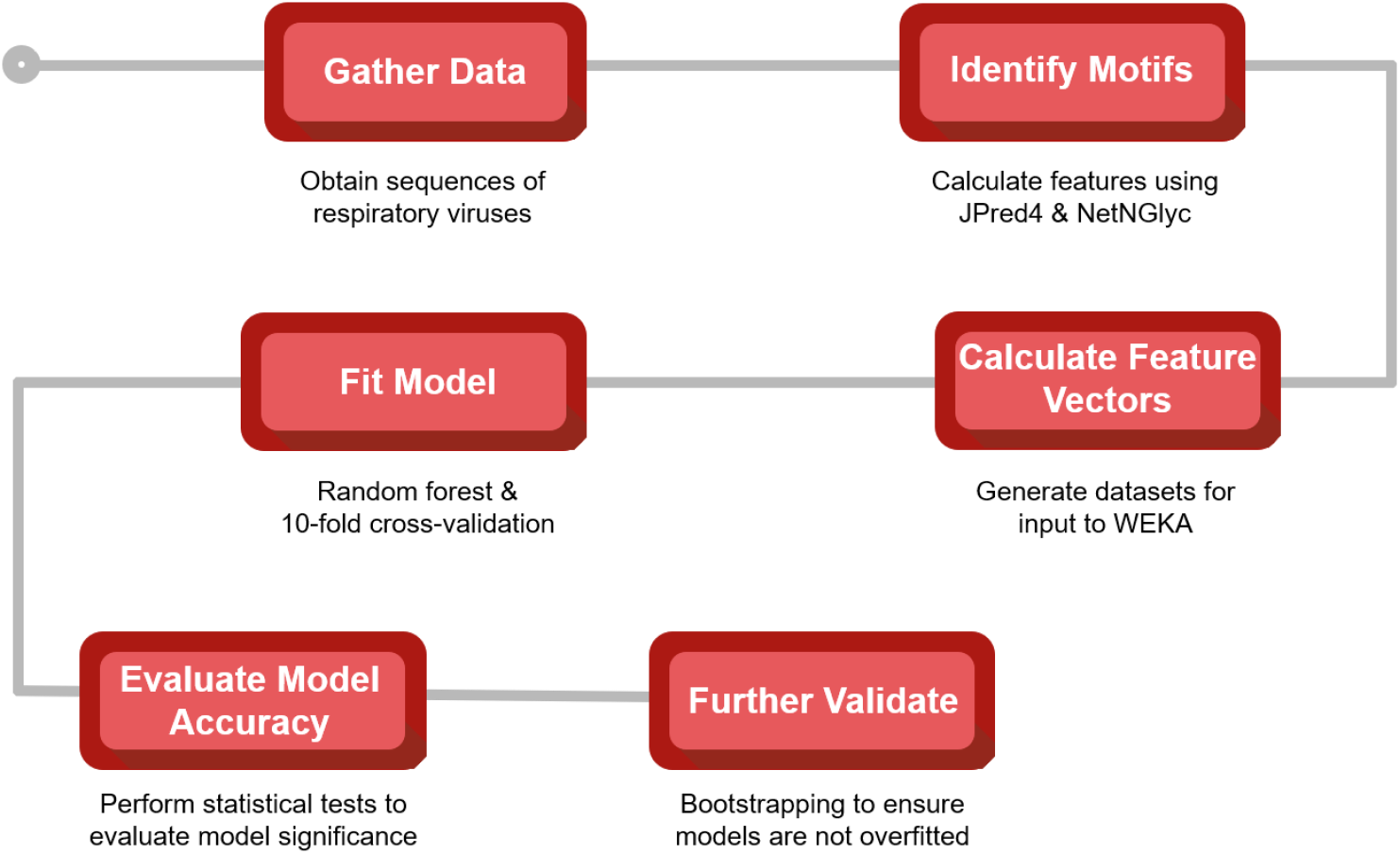
Overall model development workflow. The procedure for the development of ML models to differentiate Spike from non-Spike in a sequence.

**FIG 3.**
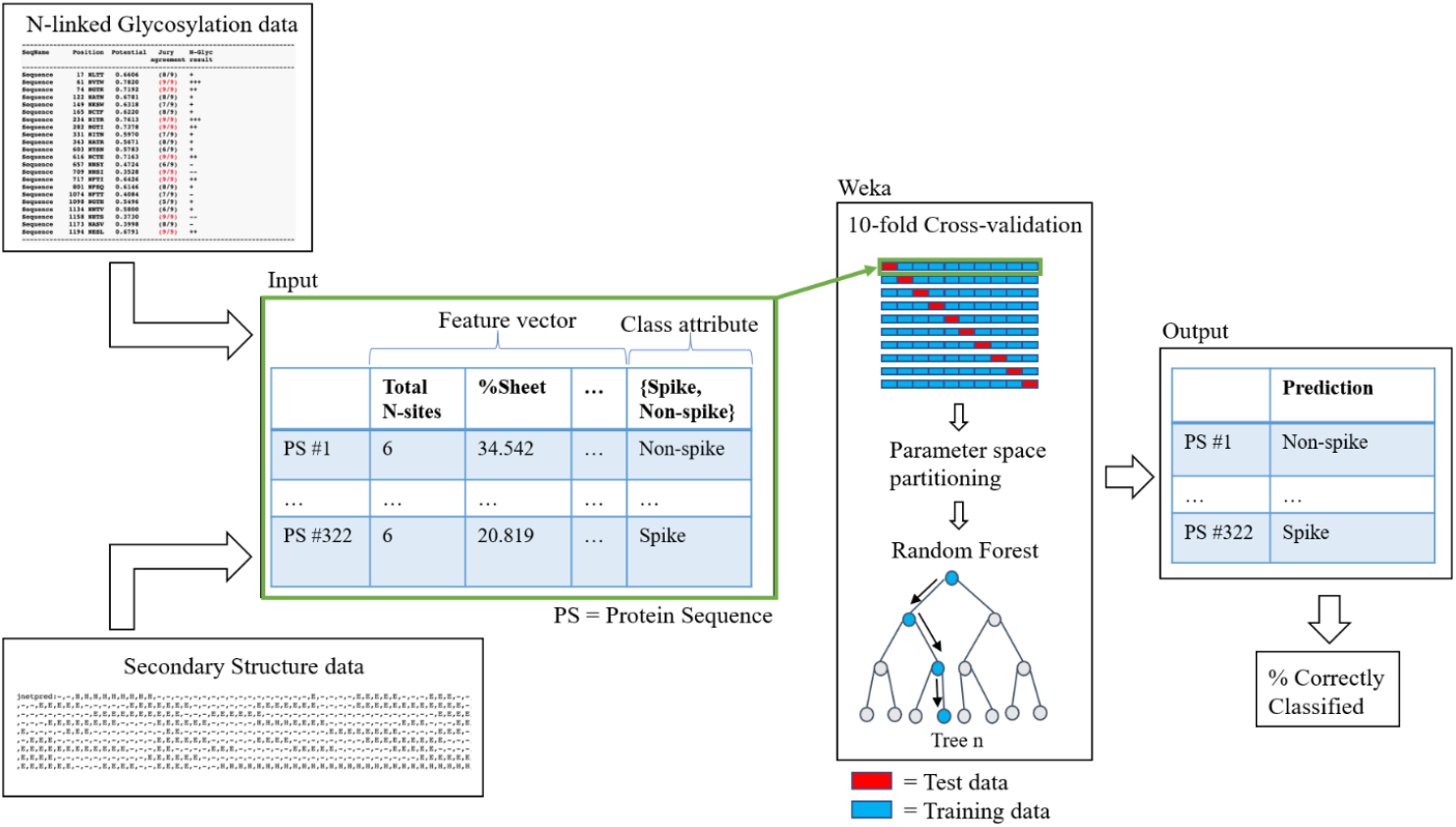
Random forest inputs, cross validation, and outputs. Data was input for 316 protein sequences.

**FIG 4.**
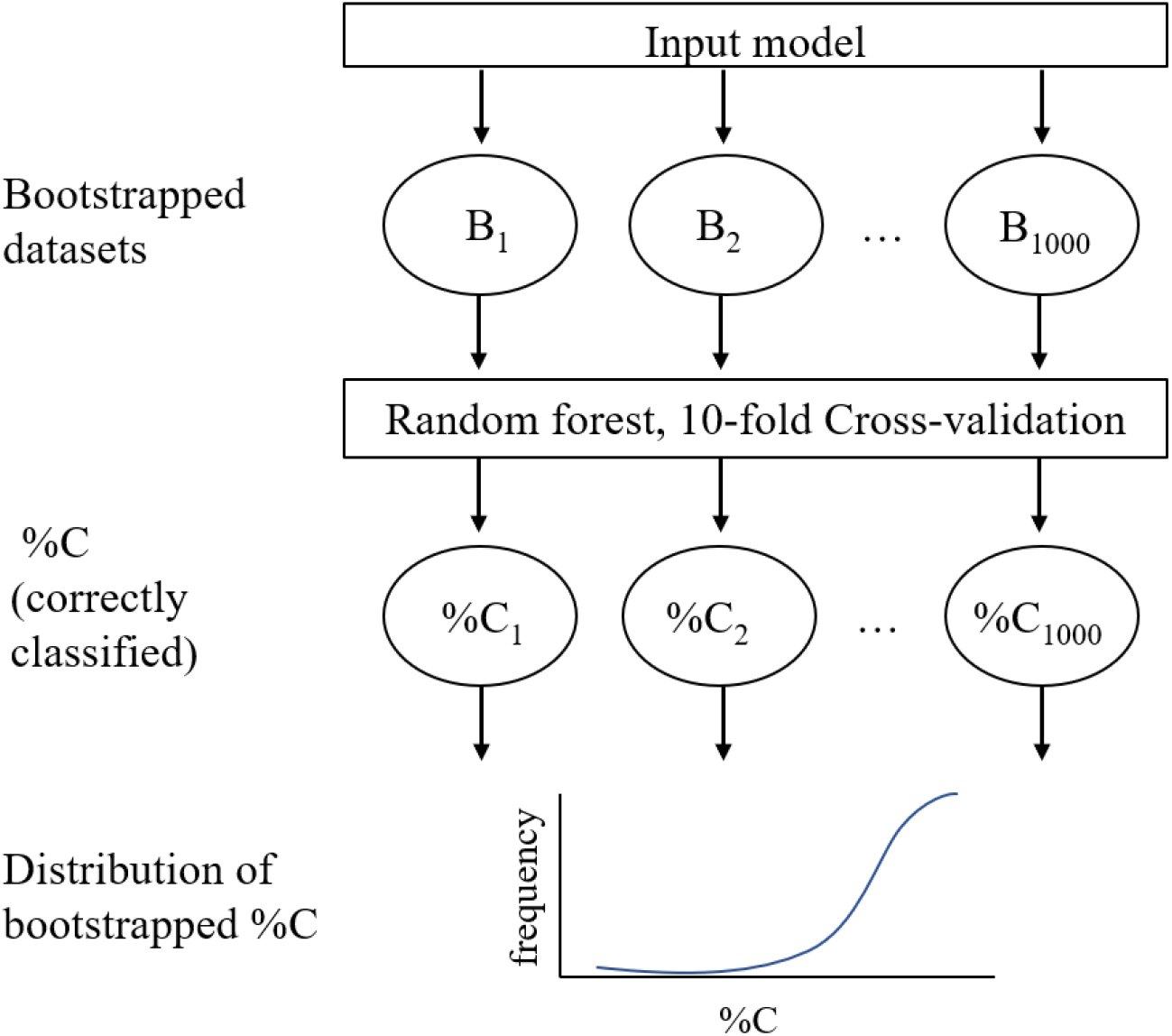
Schematic of bootstrapping process for cross validation of selected models. In this case, each of the 1000 bootstrapped datasets contains feature vectors for 316 protein sequences.

**Table 1.**
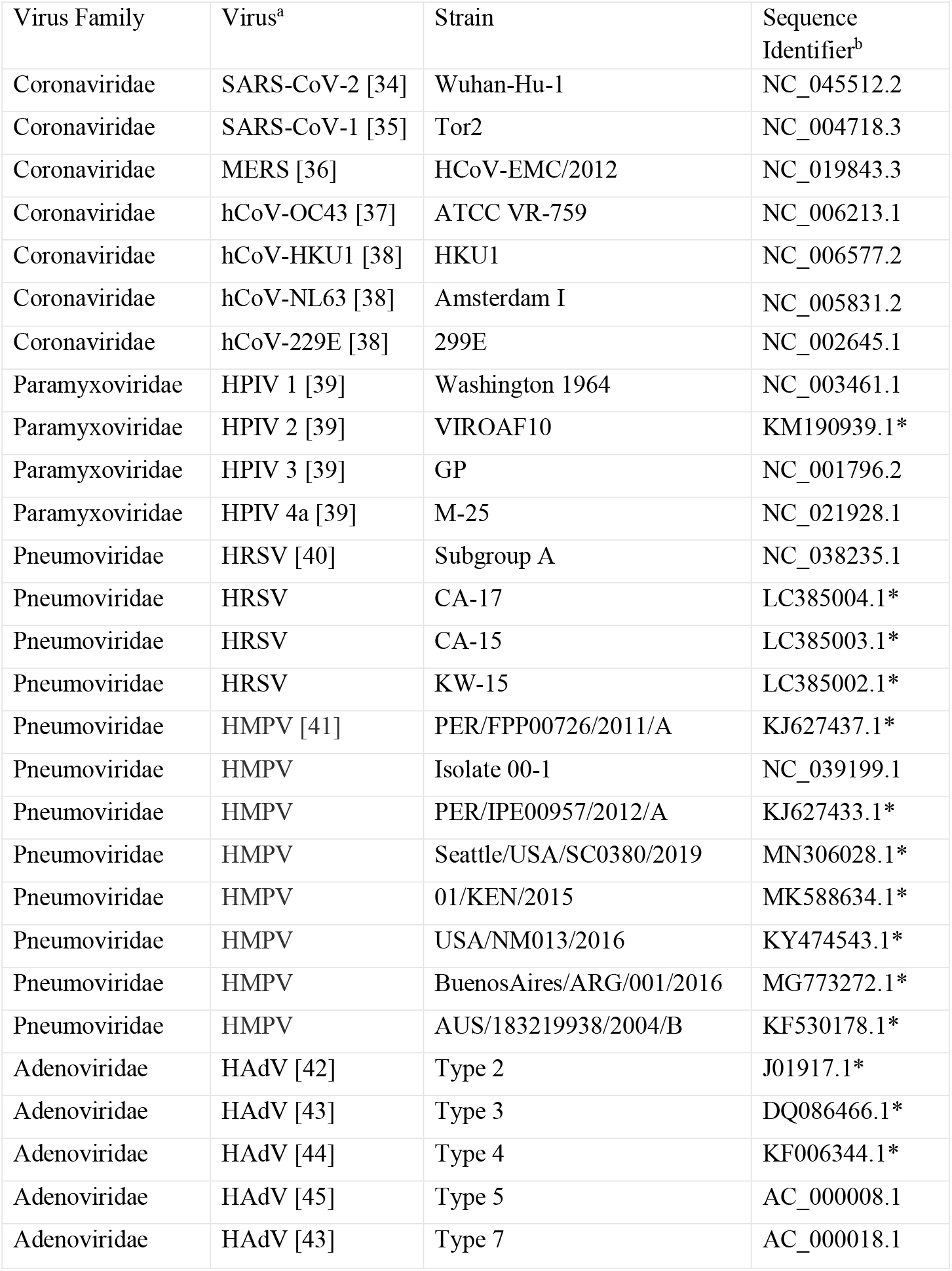
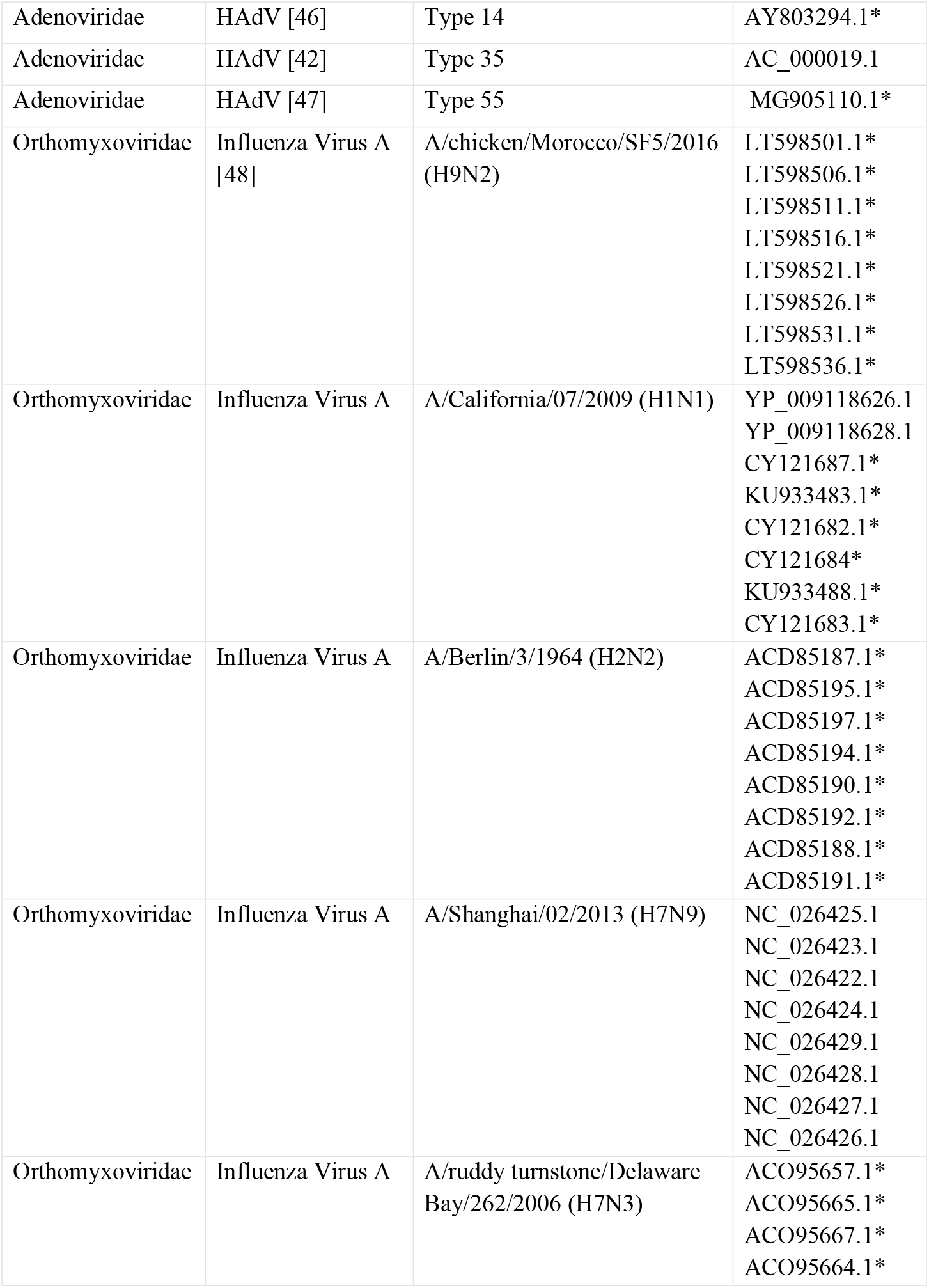
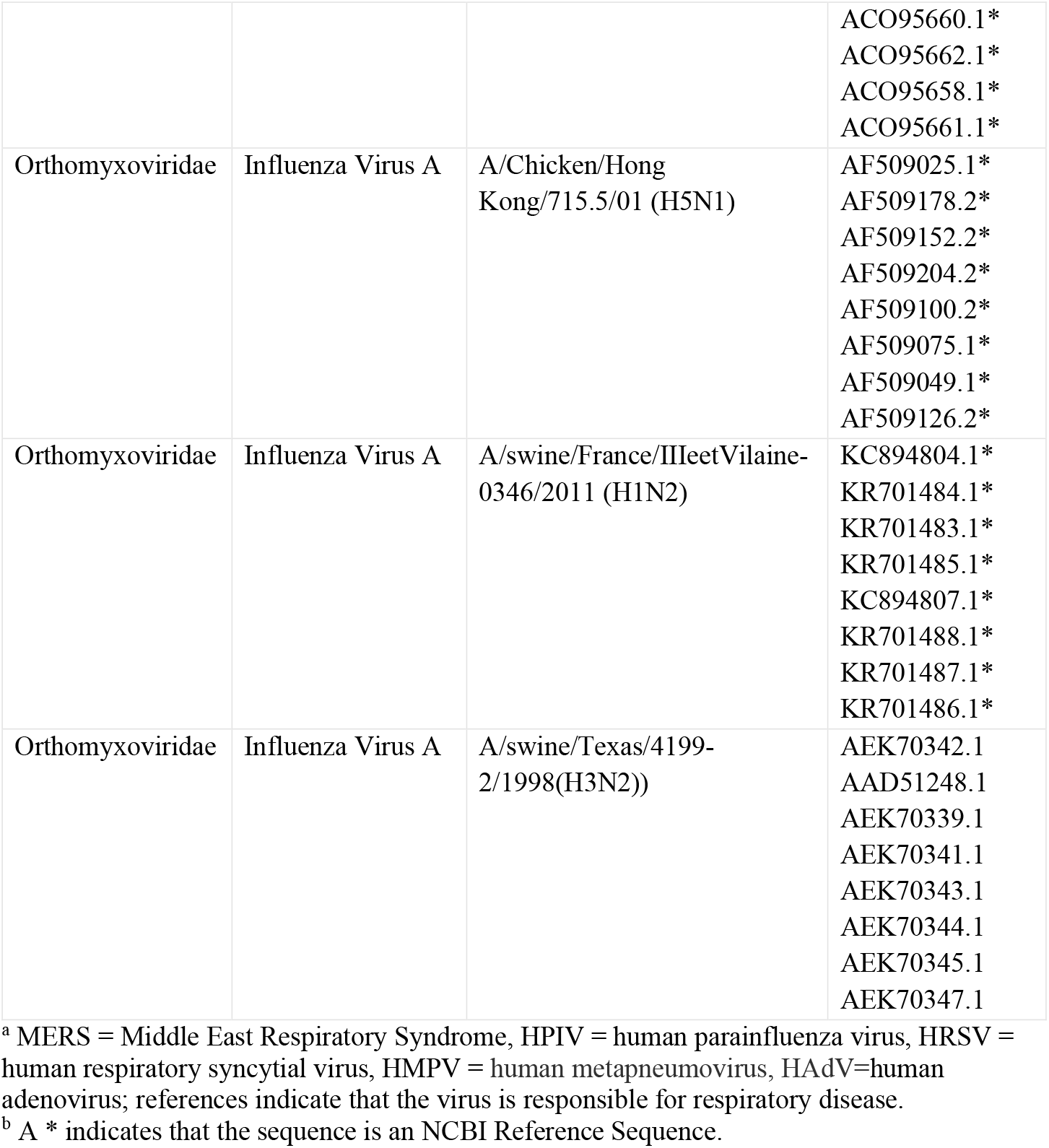
Respiratory Virus Sequences Used in Model Development.

### Prediction of secondary structural elements

The Jpred4 [25] secondary structure prediction server was used to predict structural elements for each viral sequence in the dataset. Jpred4 is a server that hosts Jnet, a neural network secondary structure prediction algorithm trained with different representations of multiple sequence alignment profiles for the same sequences [26]. Each residue in a protein sequence is designated as H (helical), E (extended sheet), or other. Since Jpred4 predicts secondary structure on protein sequences up to 800 amino acids in length, a script (Fig. S1) was written to break protein sequences into 800 residue segments and subsequently concatenated the results. For each protein, the script calculated protein length, % H, and % E, identified the longest helix and the longest sheet and calculated % longest H, and % longest E, where % longest H (E) is the length of the longest H (E) in the protein divided by the length of the protein. Finally, %helix, %sheet, %longest H, and %longest E is output.

### Prediction of N-glycosylation sites

For the sequences described above, N-glycosylation sites were predicted for each protein using NetNGlyc [27, 28]. The NetNGlyc method uses artificial neural networks to predicts N-Glycosylation sites in proteins through analysis of the sequence context of Asn-Xaa-Ser/Thr sequons. FASTA format protein sequences were entered on the NetNGlyc 1.0 Server (https://services.healthtech.dtu.dk). Asparagines with overall positive score, denoted by ‘+’, ‘++’, ‘+++’ and ‘++++’ (each counted in their respective category), where ‘++++’ indicates a prediction with highest confidence based on a combination of overall potential score and jury agreement amongst the nine neural networks utilized, were predicted to be glycosylated. The total number of glycosylation sites per protein (total N-sites) was the sum of the number of residues scored ‘+’ or higher. The density was the total sites divided by the number of residues in the protein (as reported by NetNGlyc).

### Amino Acid Composition

Protein sequences were obtained from nucleic acid sequences with Bioinformatics Toolbox in MATLAB version 2019b (MathWorks, 2021, Natick, MA, USA), and a letter frequency counter code was used to obtain the occurrence of each amino acid (AA) for each protein. The individual occurrences were divided by the corresponding protein amino acid length and multiplied by 100, giving %AA composition.

### Statistical test of association

Two-tailed t-tests for two independent samples were performed using XLSTAT v22.2.3 (Addinsoft, 2020 New York, USA) to assess the association of various features with spike vs. non-spike protein status. Features that showed a statistically significant association (*p*-value ≤ 0.05) between spike and non-spike groups and thereby rejected the null hypothesis were considered for inclusion in the ML models.

### Inputs vectors for ML models

Feature vectors were generated for each of the 316 protein sequences to create the full dataset. For each protein, the following features were calculated as described above: total N-sites, density, %M, %N, %S, %sheet, %helix, %longest sheet, and %longest helix. The designation of spike or non-spike was also included.

### Random Forest model development

Weka, an open-source software workbench for ML and data analysis [29], was utilized to develop Random Forest classifiers derived from the dataset described above. Random forest is a supervised ensemble learning method that generates a set of decision trees maximizing the separation of the classes that are sought to be discriminated [30, 31]. Subsets of the data were converted into ARFF format and uploaded to the Weka Explorer version 3.8.4 to generate specific Random Forest models (see Table S1). For each Random Forest model, a ZeroR model was also generated. Ten-fold cross-validation was utilized with both algorithms. The statistical significance of each model result was assessed by performing a Fisher’s exact test [32].

### Bootstrapping

Bootstrapping datasets were generated using the random sampling with replacement command in MATLAB version 2019b (MathWorks, 2021, Natick, MA, USA). For each model being investigated, 1000 such datasets were generated and saved as CSV files. The CSV files were converted to ARFF format using a modified csv-to-arff Python routine (obtained from github.com/anaavila). For the 50-50 balanced bootstrapping tests, for each dataset, 50% of the feature vectors were for proteins designated as spike and the other 50% were for those designated as non-spike.

## Results and Discussion

To examine the feasibility of using a machine learning model trained on viral sequences, data set was assembled consisting of 316 protein sequences for 39 respiratory viruses from five virus families, with each protein classified as “spike” (viral attachment machinery) or non-spike. Next, the associations between various features and the classification of “spike” vs. non-spike for the coronaviruses in the dataset were examined to look for signals indicating that certain feature types may help to differentiate “spike” vs. non-spike.

It has previously been shown that across coronaviruses, prior to the emergence of SARS-CoV-2, the tertiary structure of the spike protein is not conserved but the connectivity of secondary structure elements is [8]. As evidenced in Fig. 1A, the tertiary structure of the “spike” protein is clearly not conserved across different respiratory families. For the coronavirus sequences, two-tailed t-tests were performed looking at the association of %helix, %sheet, %longest sheet, %longest helix, respectively, with spike vs. non-spike status. A statistically significant association was observed for %sheet (*p*-value = 0.001), whereas none was for %helix (*p*-value = 0.087), %longest helix (*p*-value = 0.083) and %longest sheet (*p*-value = 0.208). The %longest helix was examined because when predicted secondary structure topology was examined across the SARS-CoV-2 sequence (NC_045512.2) the spike region appeared to have more longer helical segments than the other regions of the sequence; %longest sheet was added for completeness.

The pattern of N-linked glycosylation of the spike protein is highly conserved (ref) and may play a role in immune evasion [9, 33]. Again, for the coronavirus sequences, t-tests were performed examining the correlation of total N-sites and density, respectively, for spike vs. non-spike. A significant statistical difference was found for the total N-sites (*p*-value < 0.0001) and density (*p*-value = 0.010). The %AA was also examined over the coronaviruses dataset to determine if there were significant differences in amino acid composition for spike vs. non-spike. Of the 20 %AAs, a significant difference was observed for %N (*p*-value = 0.008), %S (*p*-value = 0.030), and %M (*p*-value = 0.032).

Based on these preliminary findings, we developed Random Forest machine learning classifiers with a feature vector that consisted of glycosylation, amino acid composition, and secondary structure element related features. To place these results in context, we compared classifier accuracy in each case to the ZeroR Scores for the same dataset. The ZeroR, which consists of a simple classification rule which simply predicts the majority category (class), provides a benchmark for classification performance. We also performed a test of association between the predicted and actual classes, using Fisher’s Exact Test (ref).

Our first set of Random Forest models were developed based on the coronavirus dataset (see Table S1). All but one classified the proteins correctly 100% of the time with a ZeroR Score of 86.8% and Fisher’s Exact Test of 0.006. A comparison of these five models suggests that only total N-sites and density may contribute significantly to the models. The other model (**A.1**) involving only secondary structure—%sheet, %helix, %longest sheet, %longest helix—yielded 96.2% correctly classified; that same set of features was then used to develop a model separately for each of the other four virus families. For each of these models the % correctly classified ranged from 96.2% to 100% with a sensitivity ranging from 0.86 to 1.0, and a specificity ranging from 0.98 to 1.0. To place these results in context, the ZeroR Scores for these datasets ranged from 86.8% to 88.5%. For each of the classifiers, there was a strong association between the actual classes and the class predicted by the Random Forest, with *p*-values ranging from 0.004 to 0.065. Models based on combining total N-sites, density, %sheet, %helix, and % longest helix were also generated for each virus family (**B.1**), respectively; in this case, the % correctly classified ranged from 93.5% to 100% (compared with ZeroR Scores from 86.5% to 87.5%), and Fisher’s exact test *p*-values ranging from 0.004 to 0.238. These two feature sets (associated with the A.1 and B.1 models, respectively) were then used to create cross respiratory virus family models (**A** and **B**, respectively) using the full dataset, yielding %correctly classified of 97.8% and 98.1%, respectively. A cross virus family model not including secondary structure elements (**C**) yielded significantly poorer results with a % correctly classified of 92%. These data taken together point to the robustness of the models overall.

Cross virus family models **A** and **B** are described in detail in Table 2. As a crosscheck against overfitting, we carried out bootstrapping with 1000 datasets, using resampling with replacement to generate synthetic datasets with 316 data points each. For each dataset, we built a new Random Forest model using 10-fold cross validation and evaluated its accuracy. Ninety-five percent of the Random Forest models built for the bootstrapped datasets showed an accuracy of greater than 98%, indicating that the models were in fact not overfitted to the original dataset. For model **A**, the mean and upper confidence intervals of the %correctly classified at the 95% level were 98.86% and 98.90%; while for model **B**, they were 98.82% and 98.86%.

**Table 2.**
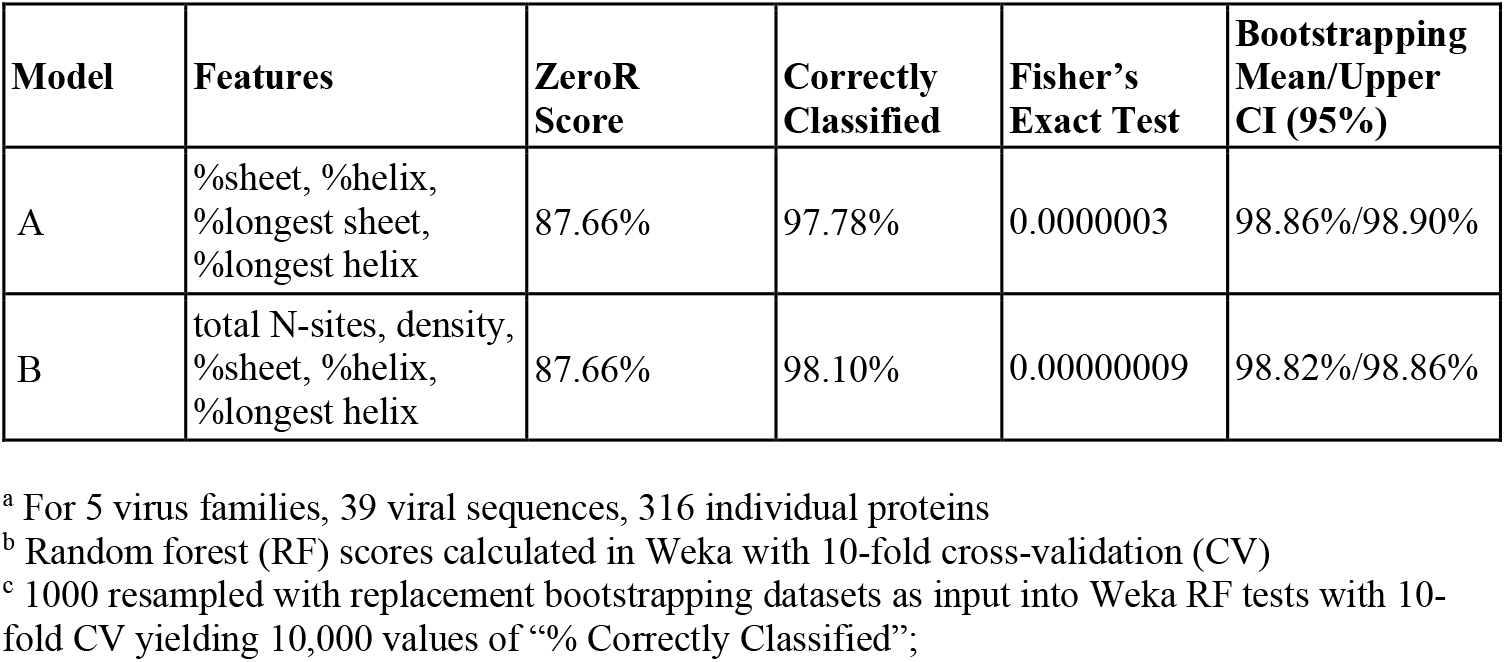
ML Models Successfully Differentiate Spike from Non-Spike^a, b, c^.

**Table 3.** ML Also Identifies Virus Families from Sequence ^a^.

| **Virus Families** | **Features** | **ZeroR Score** | **Correctly Classified** | **Fisher's Exact Test** |
| --- | --- | --- | --- | --- |
| 5 | total N-sites, density,<br>%sheet, %helix,<br>%longest helix | 32.91% | 87.97% | < 0.0000001 |

<sup>a</sup> Family class used as the output attribute (instead of “spike vs. non-spike”).

Next, bootstrapping was performed with 1000 datasets that were 50-50 balanced for “spike” vs. non-spike to eliminate the possibility that the accuracy of the models could be due to the fact that non-spike was overrepresented in the database (although obviously the proportion “spike” vs. non-spike is reflective of distribution). In this 50-50 balanced bootstrapping exercise, each dataset was comprised of 158 “spike” and 158 non-spike proteins, randomly sampled with replacement. In the balanced case, for model **A**, the mean and upper confidence intervals of the %correctly classified at the 95% level were 98.89% and 98.92%; while for model **B**, they were 99.68% and 99.84% (Table S2). This can be compared against the ZeroR score of 50% that is expected in a 50-50 balanced dataset. Thus, models **A** and **B** can successfully differentiate “spike” from non-spike respiratory virus sequence without specifying the viral family.

Finally, the capability of a ML model to identify the virus family from the sequence using the same feature vectors was explored. The Random Forest model generated was 86% accurate in predicting the virus family (see Table 2). This result is particularly impressive given that the ZeroR baseline performance indicator was only 22.36%.

In summary, the models developed by us in this work can correctly identify viral “spike” proteins from viruses (within the five viral families examined here) within seconds. The ability to utilize ML models to predict the protein responsible for cell entry (the “spike”) from a viral sequence as well as to predict the virus family of a novel viral sequence may in the future expedite the development of biomedical interventions for respiratory pandemics. In addition, the predictiveness of the models points to the underlying importance of secondary structure and N-glycosylation in viral host cell recognition.

## Acknowledgments

We would like to thank the Boston University College of Engineering STARS program for support for SD and MS.

## Supplemental Material Figure Legends

**Table S1.** All ML Models Examined.

| **Model #** | **Viral Family** | **Features** | **ZeroR Score** | **Correctly Classified** | **Fisher's Exact Test** |
| --- | --- | --- | --- | --- | --- |
| 1, **C.1** | Coronaviridae | total N-sites, density, %N, %M, %S | 86.793% | 100.000% | 0.006 |
| 2 | Coronaviridae | total N-sites, density, %M, %S | 86.793% | 100.000% | 0.006 |
| 3 | Coronaviridae | total N-sites, density | 86.793% | 100.000% | 0.006 |
| 4, **D.1** | Coronaviridae | total N-sites, density, %M, %S, %sheet, %helix, %longest sheet, %longest helix | 86.793% | 100.000% | 0.006 |
| 5, **B.1** | Coronaviridae | total N-sites, density, %sheet, %helix, %longest helix | 86.793% | 100.000% | 0.006 |
| 6, **A.1** | Coronaviridae | %sheet, %helix, %longest sheet, %longest helix | 86.793% | 96.226% | 0.065 |
| 7, **A.1** | Paramyxoviridae | %sheet, %helix, %longest sheet, %longest helix | 87.097% | 100.000% | 0.056 |
| 8, **A.1** | Pneumoviridae | %sheet, %helix, %longest sheet, %longest helix | 88.462% | 98.077% | 0.004 |
| 9, **A.1** | Adenoviridae | %sheet, %helix, %longest sheet, %longest helix | 87.500% | 98.438% | 0.015 |
| 10, **A.1** | Orthomyxoviridae | %sheet, %helix, %longest sheet, %longest helix | 87.500% | 96.875 | 0.039 |
| 11, **B.1** | Paramyxoviridae | total N-sites, density, %sheet, %helix, %longest helix | 87.097% | 93.548% | 0.238 |
| 12, **B.1** | Pneumoviridae | total N-sites, density, %sheet, %helix, %longest helix | 86.462% | 98.077% | 0.004 |
| 13, **B.1** | Adenoviridae | total N-sites, density, %sheet, %helix, %longest helix | 87.500% | 98.438% | 0.015 |
| 14, **B.1** | Orthomyxoviridae | total N-sites, density, %sheet, %helix, %longest helix | 87.500% | 96.875% | 0.039 |
| 15, **C** | Coronaviridae + Paramyxoviridae + Pneumoviridae + | total N-sites, density, %N, %M, %S | 87.658% | 92.721% | 0.011 |
|  | Adenorividae + Orthomyxoviridae |  |  |  |  |
| 16, **D** | Coronaviridae + Paramyxoviridae + Pneumoviridae + Adenorividae + Orthomyxoviridae | total N-sites, density, %M, %S, %sheet, %helix, %longest sheet, %longest helix | 87.658% | 98.101% | <0.001 |
| 17, **A** | Coronaviridae + Paramyxoviridae + Pneumoviridae + Adenorividae + Orthomyxoviridae | %sheet, %helix, %longest sheet, %longest helix | 87.658% | 97.785% | <0.001 |
| 18, **B** | Coronaviridae + Paramyxoviridae + Pneumoviridae + Adenorividae + Orthomyxoviridae | total N-sites, density, %sheet, %helix, %longest helix | 87.658% | 98.101% | <0.001 |

**Table S2.** ML Models Successfully Differentiate Spike from Non-Spike ^a, b, c^ for a 50-50 balanced bootstrapping dataset ^d^.

| **Model\*** | **Features** | **Bootstrapping**<br>**Mean/Upper CI**<br>**(95%)** |
| --- | --- | --- |
| A-50-50 | %sheet, %helix,<br>%longest sheet,<br>%longest helix | 98.89% / 98.92% |
| B-50-50 | total N-sites, density,<br>%sheet, %helix,<br>%longest helix | 99.68%/99.84% |

<sup>a</sup> For 5 virus families, 39 viral sequences, 316 individual proteins
<sup>b</sup> Random forest (RF) scores calculated in Weka with 10-fold cross-validation (CV)
<sup>c</sup> 1000 resampled with replacement bootstrapping datasets as input into Weka RF tests with 10-fold CV yielding 10,000 values of “% Correctly Classified”
<sup>d</sup> Each model has 158 spike and 158 non-spike proteins, randomly sampled with replacement

**FIG S1:**
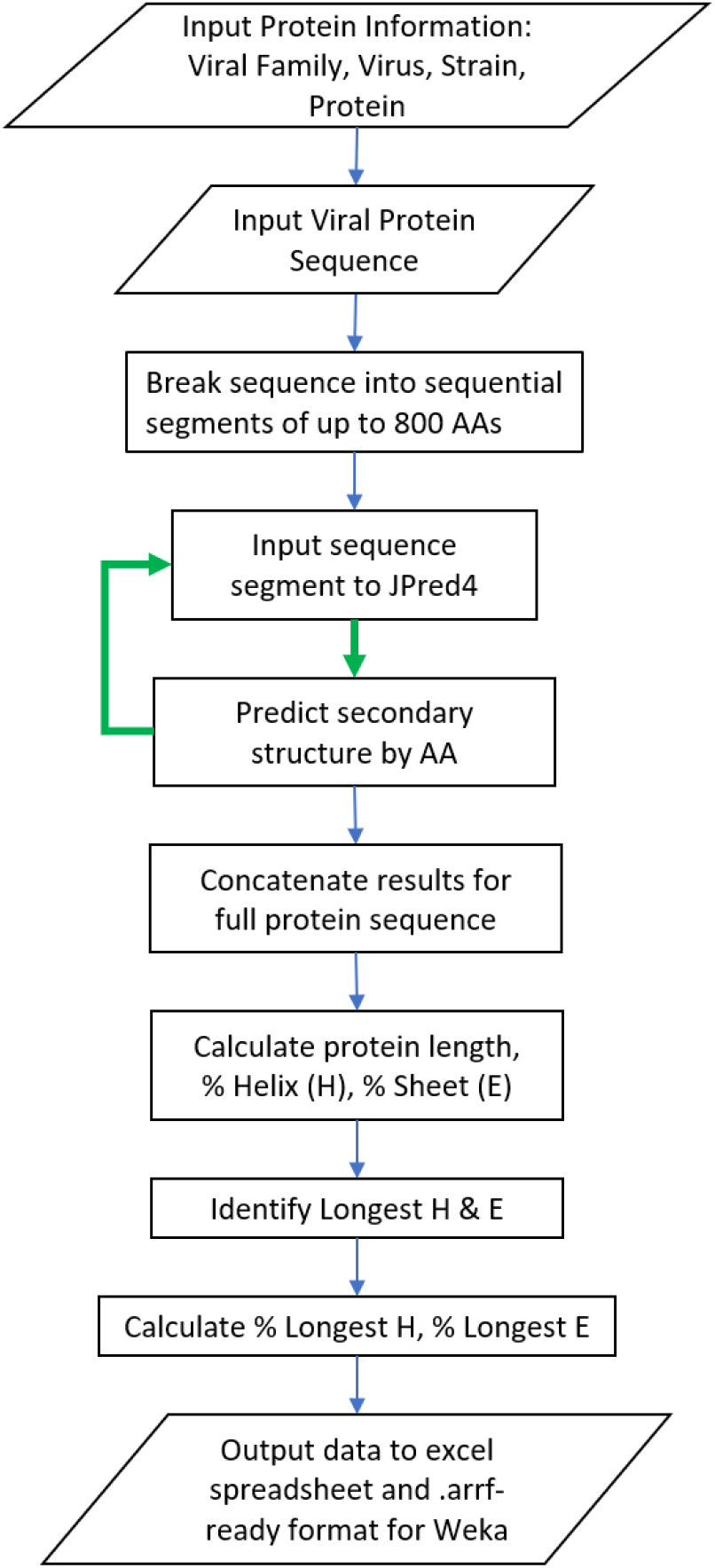
Calculation of secondary structure elements. A flowchart showing the process for calculating the secondary structure elements with Jpred4.

